## Supplementary figures for "A new mechanism of posttranslational polyglutamylation regulates phase separation and signaling of the Wnt pathway protein Dishevelled"

**This PDF file includes:**

Figs. S1 to S10
Tables S1 to S5

Fig. S1

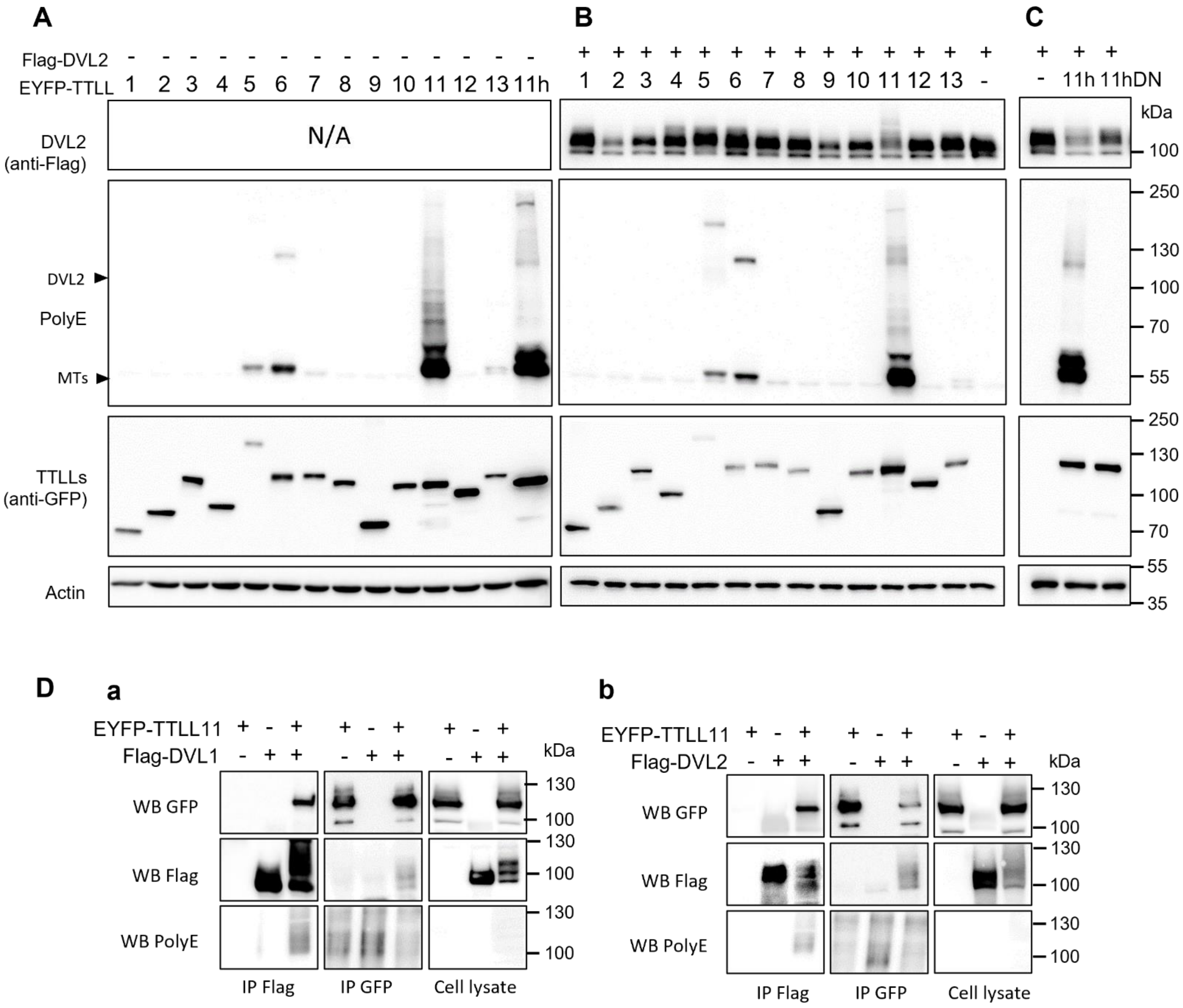


**Fig. S1. TTLL11 binds and polyglutamylates DVL1 and DVL2.**

(**A**-**C**) HEK293T cells were transfected with constructs encoding murine YFP tagged TTLL1 – TTLL13 together with control plasmid (A) or DVL2 (B), or DVL2 with human TTLL11 and its inactive variant TTLL11 (E531G; DN) (C). The samples were subjected to WB analysis using polyglutamylation specific antibody – PolyE. Note the appearance of polyE-positive bands of DVL2 size in conditions where TTLL11 and DVL2 were co-expressed. (**D**) Co-immunoprecipitation of Flag-DVL1 (**a**) or Flag-DVL2 (**b**) with TTLL11 overexpressed in HEK293T cells. TTLL11 was co-immunoprecipitated with both DVL1 and DVL2 and both DVLs were polyglutamylated in the pulldown when co-expressed with TTLL11 (IP Flag, WB PolyE).

### Fig. S2

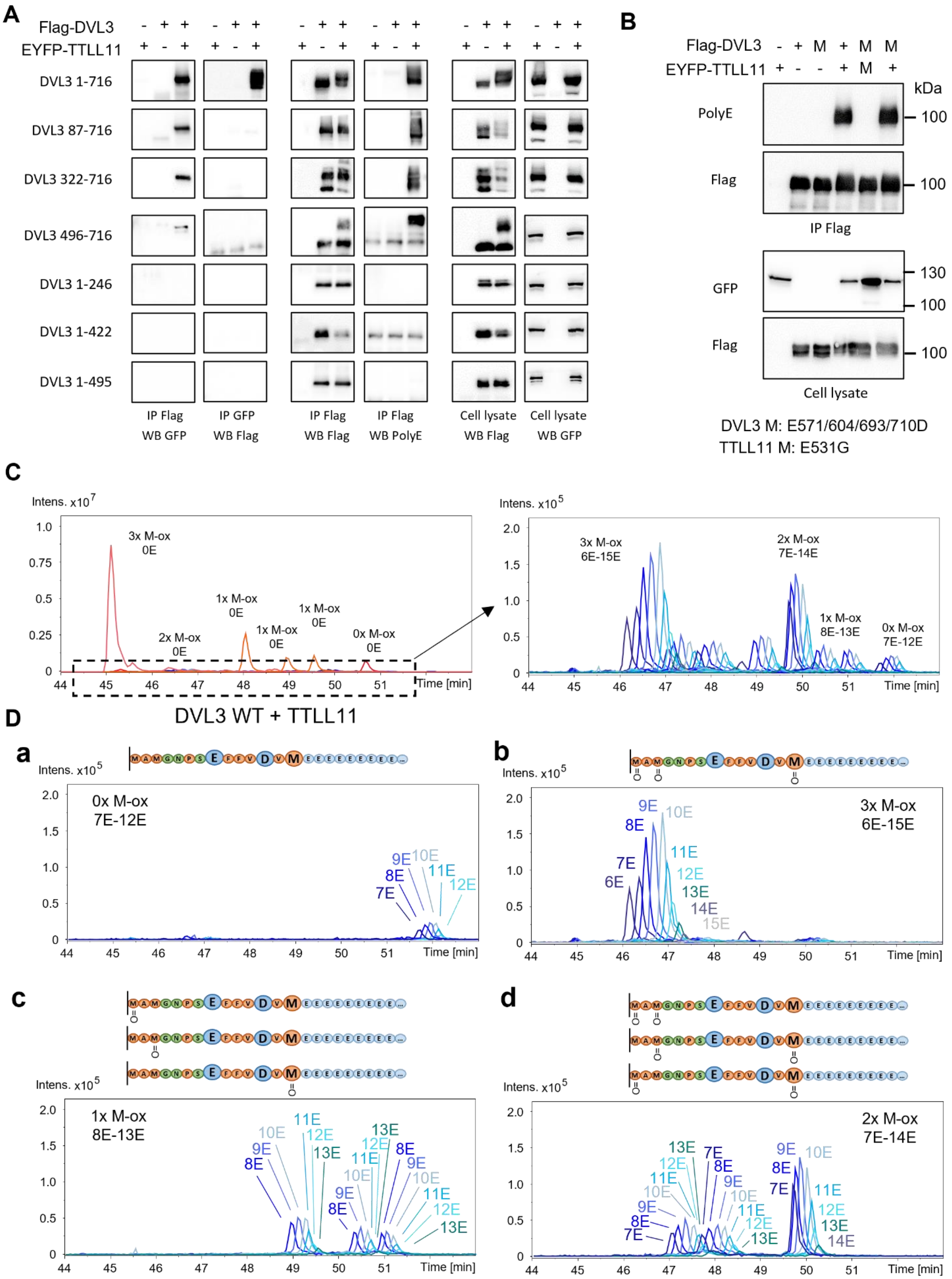
A
Flag-DVL3
EYFP-TTLL11
DVL3 1-716
DVL3 87-716
DVL3 322-716
DVL3 496-716
DVL3 1-246
DVL3 1-422
DVL3 1-495
IP Flag WB GFP
IP GFP WB Flag
IP Flag WB Flag
IP Flag WB PolyE
Cell lysate WB Flag
Cell lysate WB GFP
B
Flag-DVL3
EYFP-TTLL11
kDa
PolyE
Flag
IP Flag
GFP
Flag
Cell lysate
DVL3 M: E571/604/693/710D
TTLL11 M: E531G
C
Intens. x10^7
3x M-ox 0E
2x M-ox 0E
1x M-ox 0E
0x M-ox 0E
Time [min]
Intens. x10^5
3x M-ox 6E-15E
2x M-ox 7E-14E
1x M-ox 8E-13E
0x M-ox 7E-12E
DVL3 WT + TTLL11
D
a
0x M-ox 7E-12E
b
3x M-ox 6E-15E
c
1x M-ox 8E-13E
d
2x M-ox 7E-14E

**Fig. S2. Modification site of DVL3 polyglutamylation.**

(**A**) Domain mapping of DVL3 polyglutamylation and search for TTLL11 interaction domain on DVL3. DVL3 and its truncated mutants were co-expressed with TTLL11 in HEK293T cells and DVL3 was subsequently immunoprecipitated. Only the DVL3 mutants containing C-terminal part were able to pull-down TTLL11. Samples were also stained by modification-specific antibody PolyE, which detected polyglutamylation in all DVL3 mutants containing its C-terminus. (**B**) Polyglutamylation of DVL3 E571D/E604D/E693D/E710D is comparable to DVL3 wt. DVL3 variants were overexpressed with TTLL11 or with its inactive mutant TTLL11 E531G in HEK293 cells and immunoprecipitated via Flag tag and analysed by WB as indicated (**C**, **D**) EIC chromatogram shows peaks corresponding to the very last C-terminal peptide of DVL3 formed after tryptic cleavage and its polyglutamylated variants, that are highlighted in a separate window. The data are from the same experiment as Fig S3A. (C) EIC shows peaks corresponding to polyglutamylated peptides shown schematically above each chromatogram. (D) chromatograms for non- (D**a**), mono- (D**c**), di- (D**d**) and tri-oxidized (D**b**) peptides are presented separately.

Fig. S3

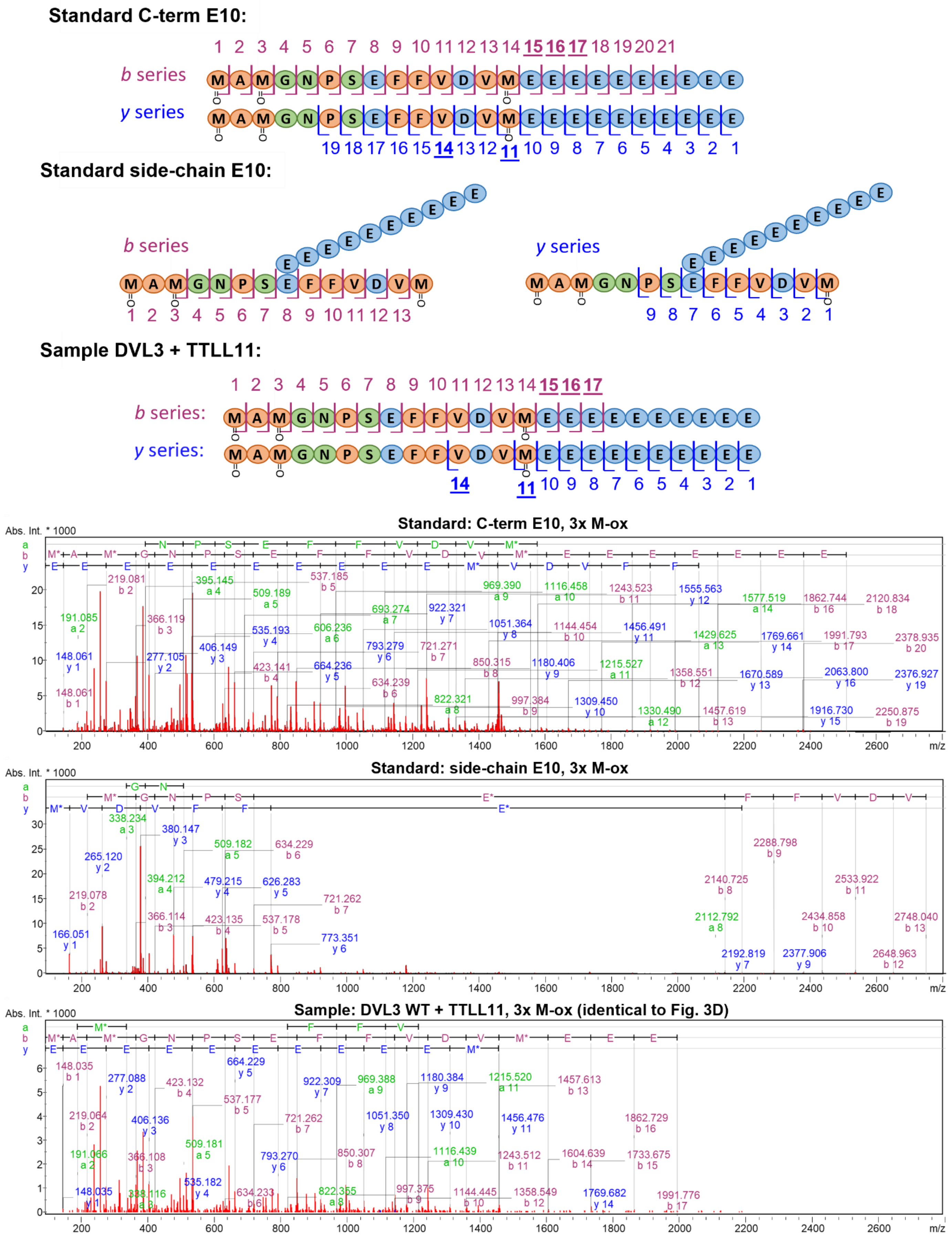

Standard C-term E10:
b series
y series
Standard side-chain E10:
b series
y series
Sample DVL3 + TTLL11:
b series:
y series:
Standard: C-term E10, 3x M-ox
Standard: side-chain E10, 3x M-ox
Sample: DVL3 WT + TTLL11, 3x M-ox (identical to Fig. 3D)
Abs. Int. * 1000
m/z

**Fig. S3. A novel character of DVL3 PTM – MS/MS spectra.**

MS/MS spectra of control synthetic peptides with 10E modification at M716 or E710 in comparison to the polyglutamylated DVL3 immunoprecipitated from cells. The aa sequences for individual peaks a shown in schematic representation above individual peaks in B and Y series. The fragments corresponding to modification on E710 are detected only in synthetic peptide control but not in DVL3 WT sample.

Fig. S4

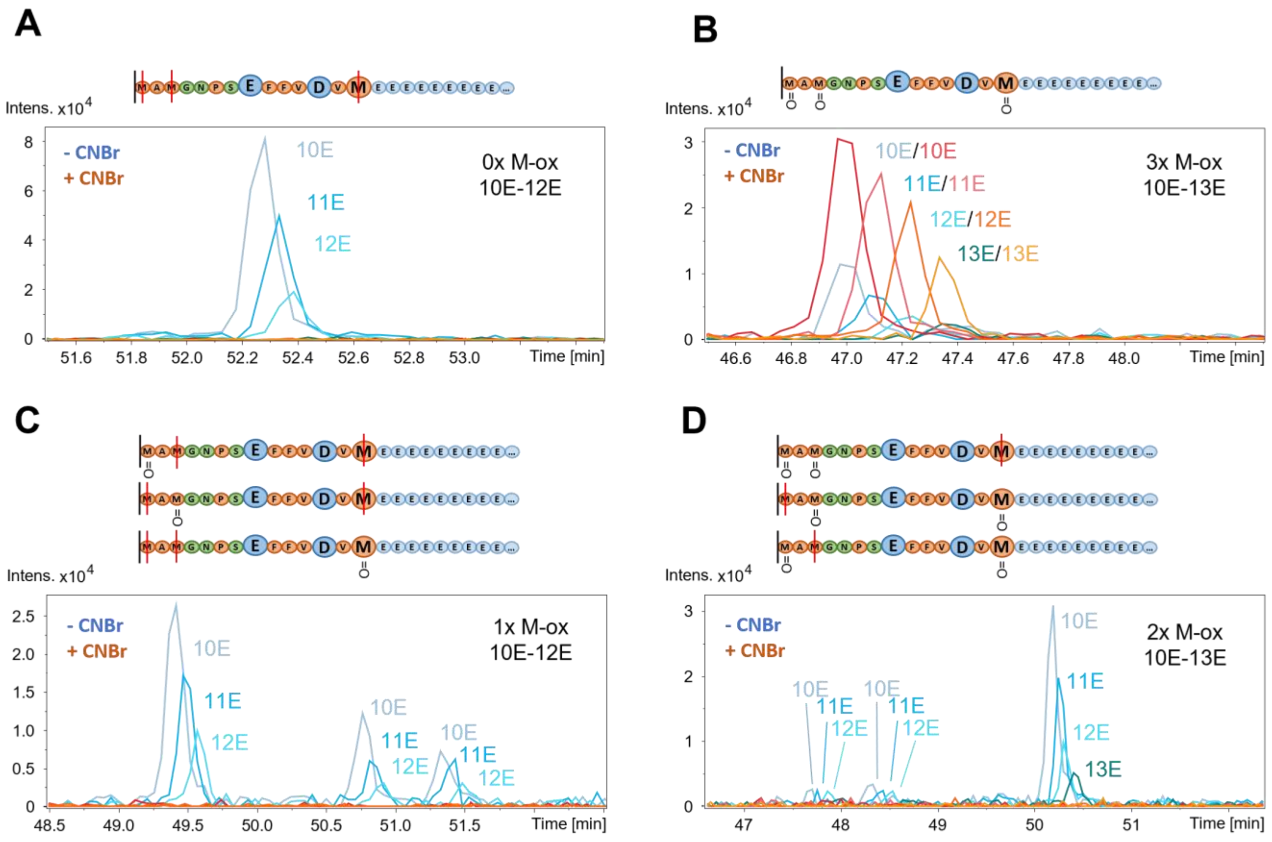


**Fig. S4. A novel character of DVL3 PTM – CNBr cleavage.**

(**A-D**) CNBr cleavage of polyglutamylated DVL3 C-terminal peptide in all its M-ox forms. EIC shows peaks corresponding to polyglutamylated peptides shown schematically above each chromatogram. Red line indicates cleavage site. A, C and D shows 0x, 1x and 2x M-ox peptides respectively, where all peptides were cleaved by CNBr. B chromatogram shows 3x M-ox peptide that cannot be cleaved by CNBr.

Fig. S5

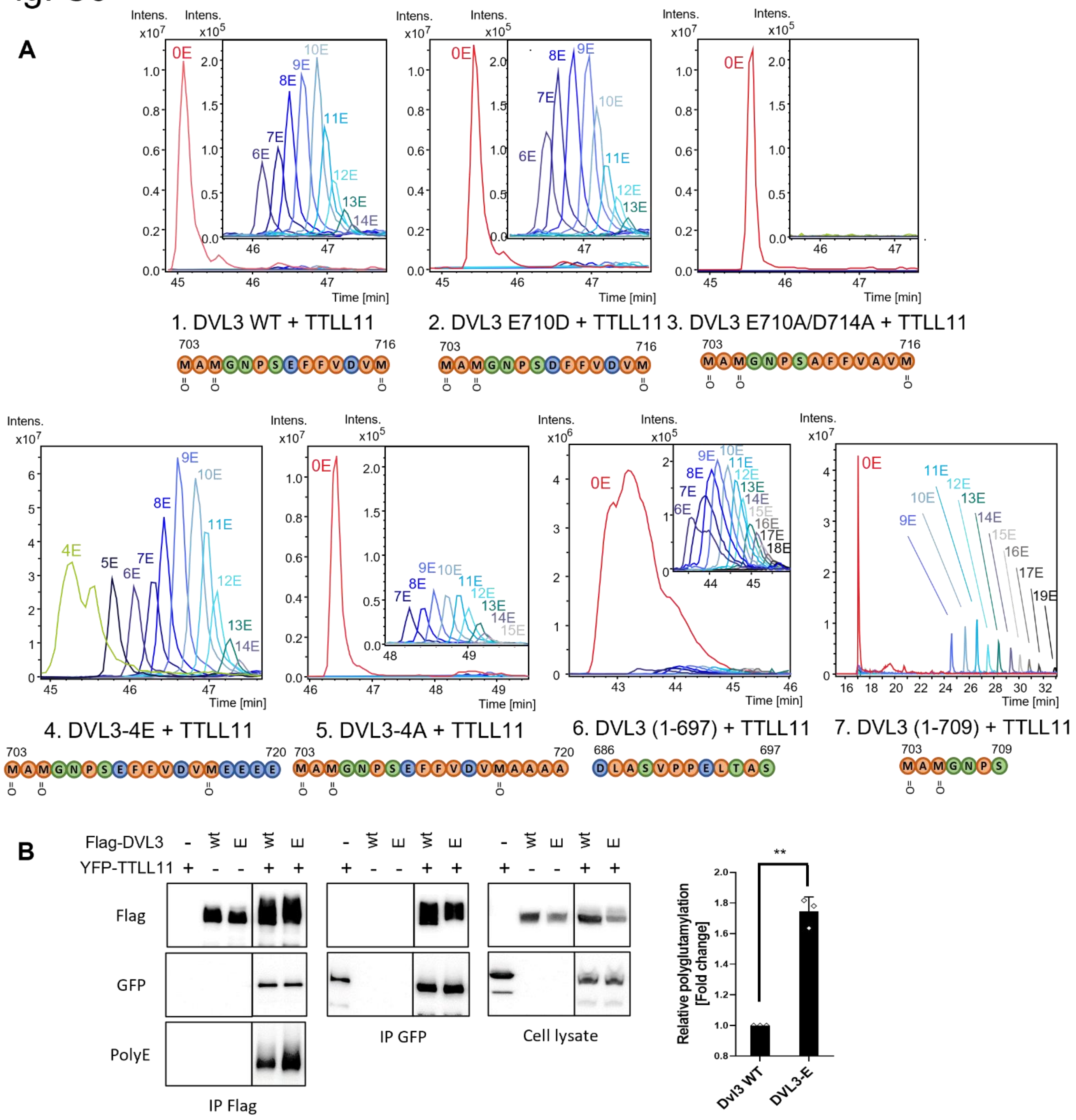


**Fig. S5. C-terminal polyglutamylation.**

(**A**) EIC showing polyglutamylation of DVL3 C-terminus by TTLL11 in WT protein and mutated variants from Fig 3A. Lower intensity peaks for polyglutamylated peptides are highlighted in separate windows. Results are shown for fully M-ox peptides. (**B**) Polyglutamylation of DVL3 C-terminus after addition of 1E residue at the C-terminal M716. Relative polyglutamylation was derived from PolyE band intensity, normalized to total protein (Flag) signal, and is shown as a fold change compared to Dvl3 WT polyglutamylation in three independent replicates. Statistical significance was assessed using one-sample t-test; **represents $p < 0.01$.

Fig. S6

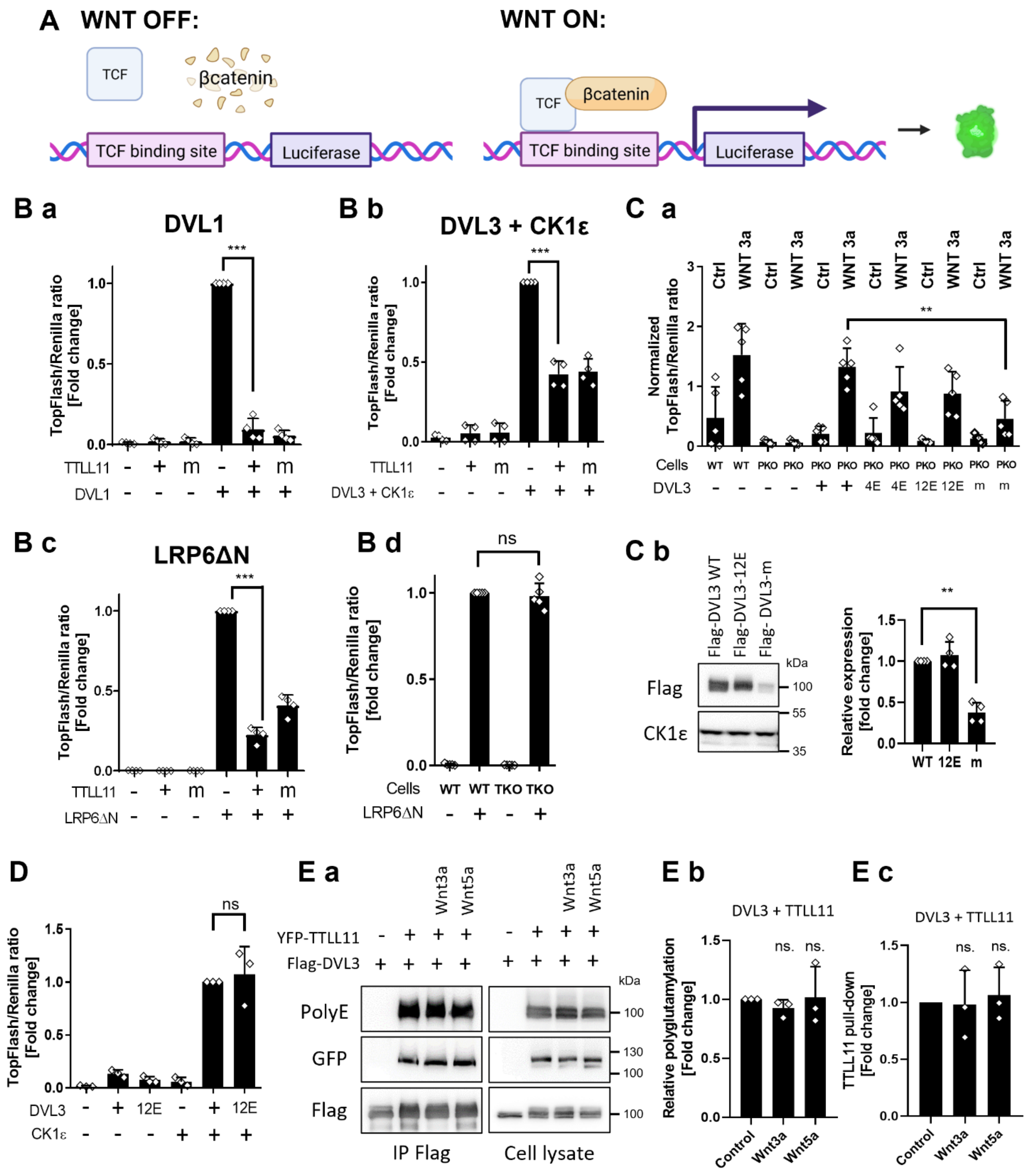


**Fig. S6. DVL3 polyglutamylation does not affect activity in the Wnt/β-catenin pathway.**

(**A**) TopFlash scheme created with BioRender.com. WNT/β-catenin pathway activation results in the production of active luciferase. (**B**) TopFlash reporter assay of TTLL11 and its inactive mutant E531G (indicated as m) with Wnt/β-catenin pathway activators: DVL1 (B**a**), DVL3+CK1 (B**b**) and LRP6ΔN (B**c**). (B**d**) TopFlash reporter assay of LRP6ΔN in Hek293 T-rex (WT) or HEK293 T-rex

DVL1/TVL2/DVL3 KO cells (TKO). The data are represented as fold change compared to the inducer sample. (**C**) DVL3 rescue assays in HEK293 T-rex RNF43/ZNRF3/DVL1/DVL2/DVL3 penta knockout cells (PKO) that lack endogenous DVL and cannot respond to Wnt-3a. (C**a**) PKO cells were transfected by indicated DVL3 variant and Wnt/β-catenin pathway signaling was induced by Wnt3a. WT cells = control. (C**b**) DVL3m, DVL3-WT and DVL3-12E expression in cells transfected by the same amount of DNA. WB intensities are normalized to DVL3-WT; n=4. CK1ε = loading control. (**D**) Both DVL3 and DVL3-12E potently activate TopFlash assay upon CK1ε co-expression. (**Ea**) Analysis of DVL3 polyglutamylation and interaction with TTLL11 after stimulation by Wnt-3a or Wnt-5a. WB intensities for polyglutamylation (PolyE; E**b**) or TTLL11 co-purification (GFP; E**c**) were normalized to Flag signal (DVL3 amount). Results from 3 replicates are shown as a fold change to control polyglutamylation (b) or pull-down (c). TopFlash data represent mean ± SD; n=4 for B; n=5 for C; n=3 for D. ** represents $p < 0.01$, *** $p < 0.001$, ns = not significant. Statistics: one-sample t-test for B, Cb, D, Eb and E; one-way ANOVA with Dunnett` multiple comparisons test for Ca. (B, D and E) were performed in HEK293T cells.

### Fig. S7

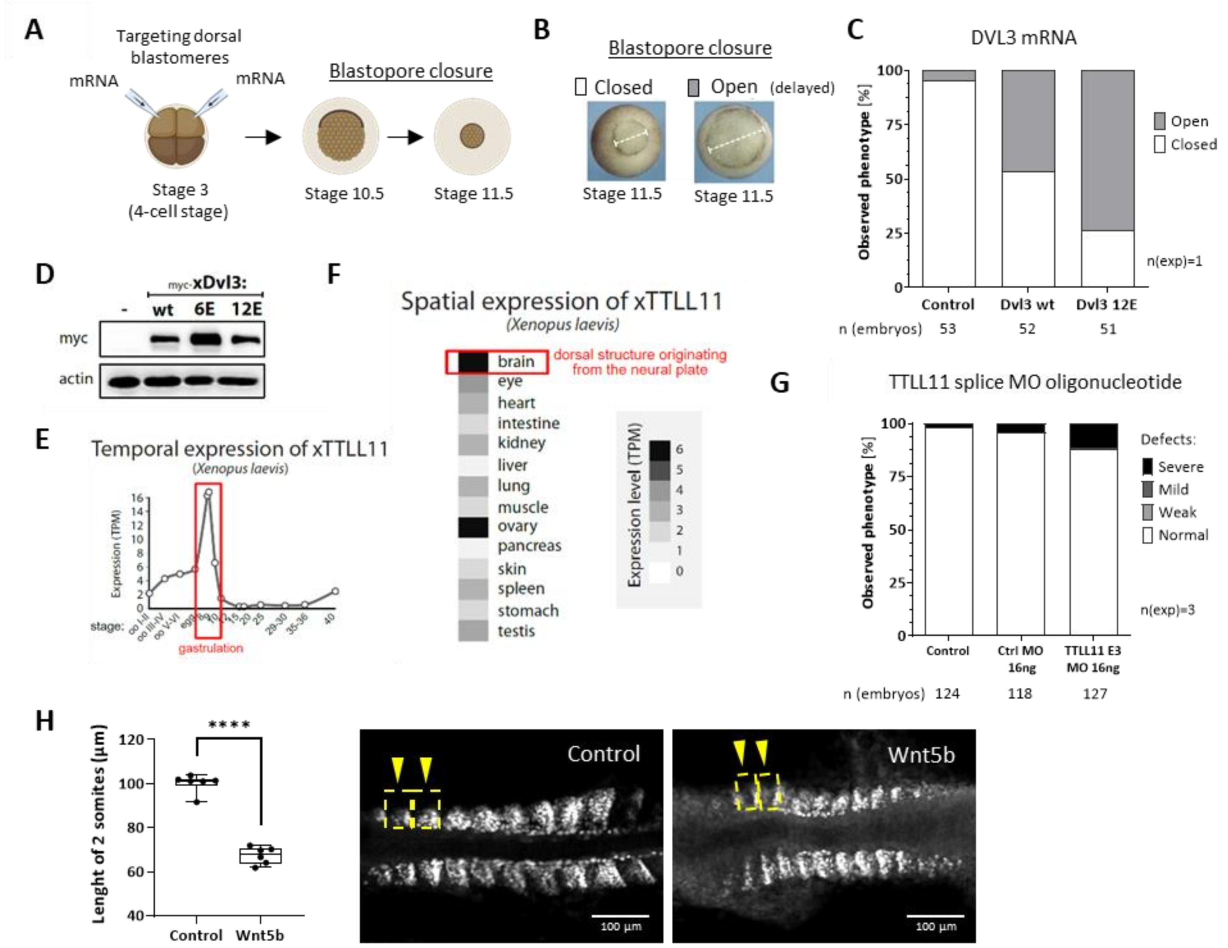


**Figure S7. Role of DVL3 polyglutamylation in *Xenopus laevis* and *Danio rerio* embryonal development.** (**A**) Both dorsal blastomeres of 4-cell *Xenopus laevis* embryo were injected with xDvl3 mRNA and embryos were observed during blastopore closure at stage 11.5 (scheme created with BioRender.com). (**B**) Normal or delayed blastopore closure was assessed at stage 11.5. (**C**) Effect on uninjected embryos and embryos injected with either mRNA of xDvl3 WT and xDvl3-12E modification mimicking mutant. (**D**) WB analysis of xDvl3 protein amount in *Xenopus* embryo lysates. (**E**) xTTLL11 RNA expression during the development of *Xenopus laevis*. (**F**) Spatial expression of xTTLL11 in adult organism (E, F adapted from (79)). (G) Effect of splicing MO targeting xTTLL11 exon 3 (Suppl. To Fig. 5D). (**H**) The length of the first 2 anterior somites was measured for control embryo Wnt5b KO embryos (n=6). Statistical analysis was performed by unpaired t test.

Fig. S8

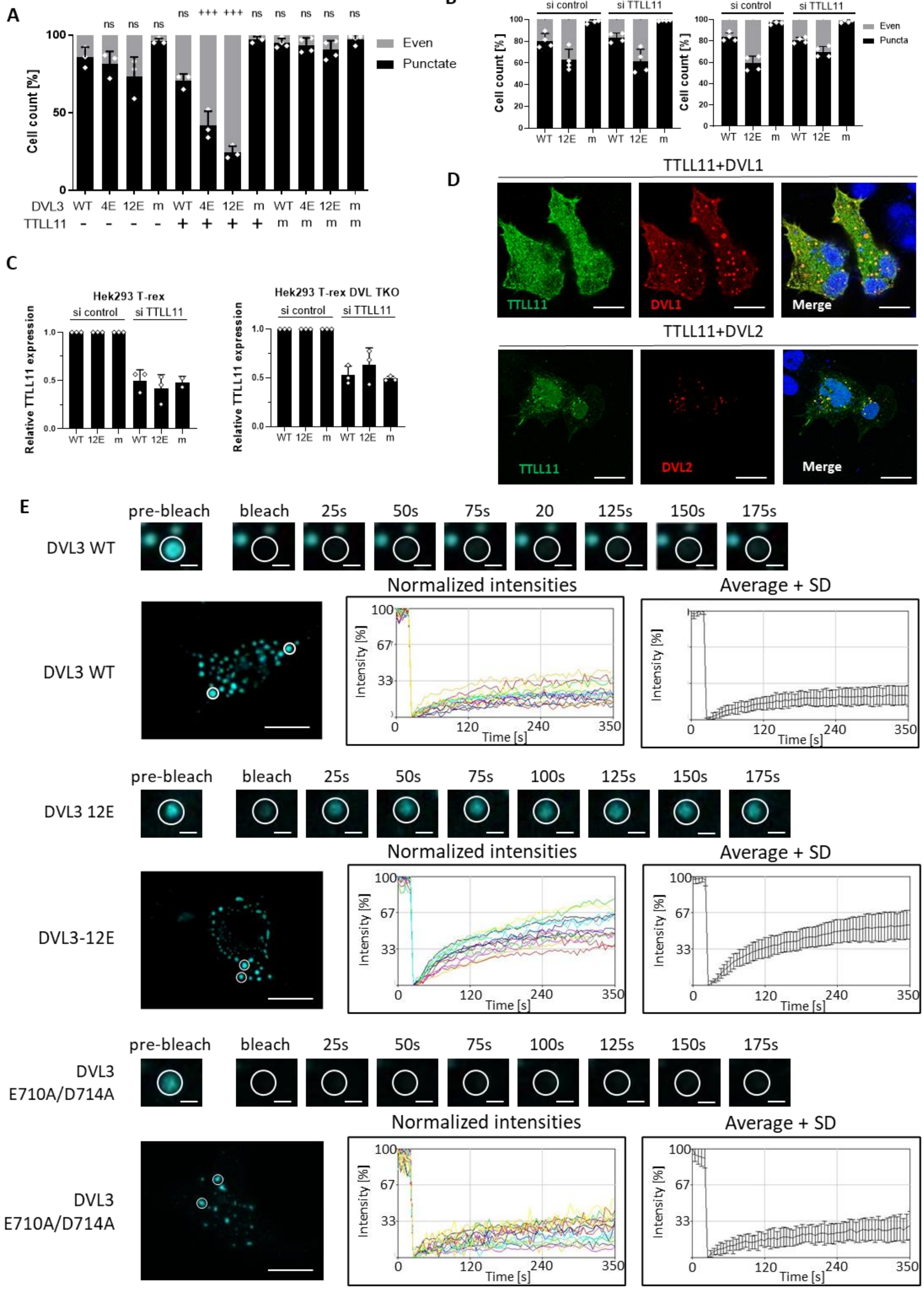
A
Cell count [%]
Even
Punctate
ns ns ns +++ +++ ns ns ns ns ns
DVL3 WT 4E 12E m WT 4E 12E m WT 4E 12E m
TTLL11 - - - - + + + + m m m m
B
Hek293 T-rex
Hek293 T-rex DVL TKO
si control
si TTLL11
Cell count [%]
Even
Puncta
WT 12E m WT 12E m
C
Hek293 T-rex
Hek293 T-rex DVL TKO
si control
si TTLL11
Relative TTLL11 expression
WT 12E m WT 12E m
D
TTLL11+DVL1
TTLL11
DVL1
Merge
TTLL11+DVL2
TTLL11
DVL2
Merge
E
pre-bleach
bleach
25s
50s
75s
20
125s
150s
175s
DVL3 WT
Normalized intensities
Average + SD
Intensity [%]
Time [s]
pre-bleach
bleach
25s
50s
75s
100s
125s
150s
175s
DVL3 12E
DVL3-12E
Normalized intensities
Average + SD
pre-bleach
bleach
25s
50s
75s
100s
125s
150s
175s
DVL3 E710A/D714A
Normalized intensities
Average + SD

**Figure S8. The effects of DVL3 polyglutamylation on the DVL3 LLPS.** Analysis of punctate vs. even subcellular localization of overexpressed DVL3 and its polyglutamylated variant or DVL3m = DVL3 E710A/D714A in (**A**) with co-expressed TTLL11 in HEK293T cells or (**B**) with or without TTLL11 downregulation by siRNA in HEK293T T-rex WT or HEK293T T-rex DVL1-3 triple KO (TKO) cells. DVL3 localization (A, B) was assessed for at least 200 cells in at least three independent replicates. SD and individual data points (white dots) are indicated. Statistically significant difference from WT DVL3 (* - all not significant, n.s.) and from the same DVL variant in the absence of TTLL11 (+) is indicated. (**C**) Quantitative PCR analysis of TTLL11 expression (Normalized on Rps13 housekeeping gene) in the samples from B. (**D**) Co -localization of overexpressed DVL1 and DVL2 with TTLL11 in HEK293T cells. TTLL11 co-localizes with DVL1 and DVL2 in puncta. Scale bar = 10 µm. (**E**) FRAP analysis of DVL3 wt, DVL3-12E and DVL3 E710A/D714A intracellular condensates. The data represent normalized intensities and average + SD from 15 condensates measured in 3 biological replicates (5 condensates per replicate).

Fig. S9

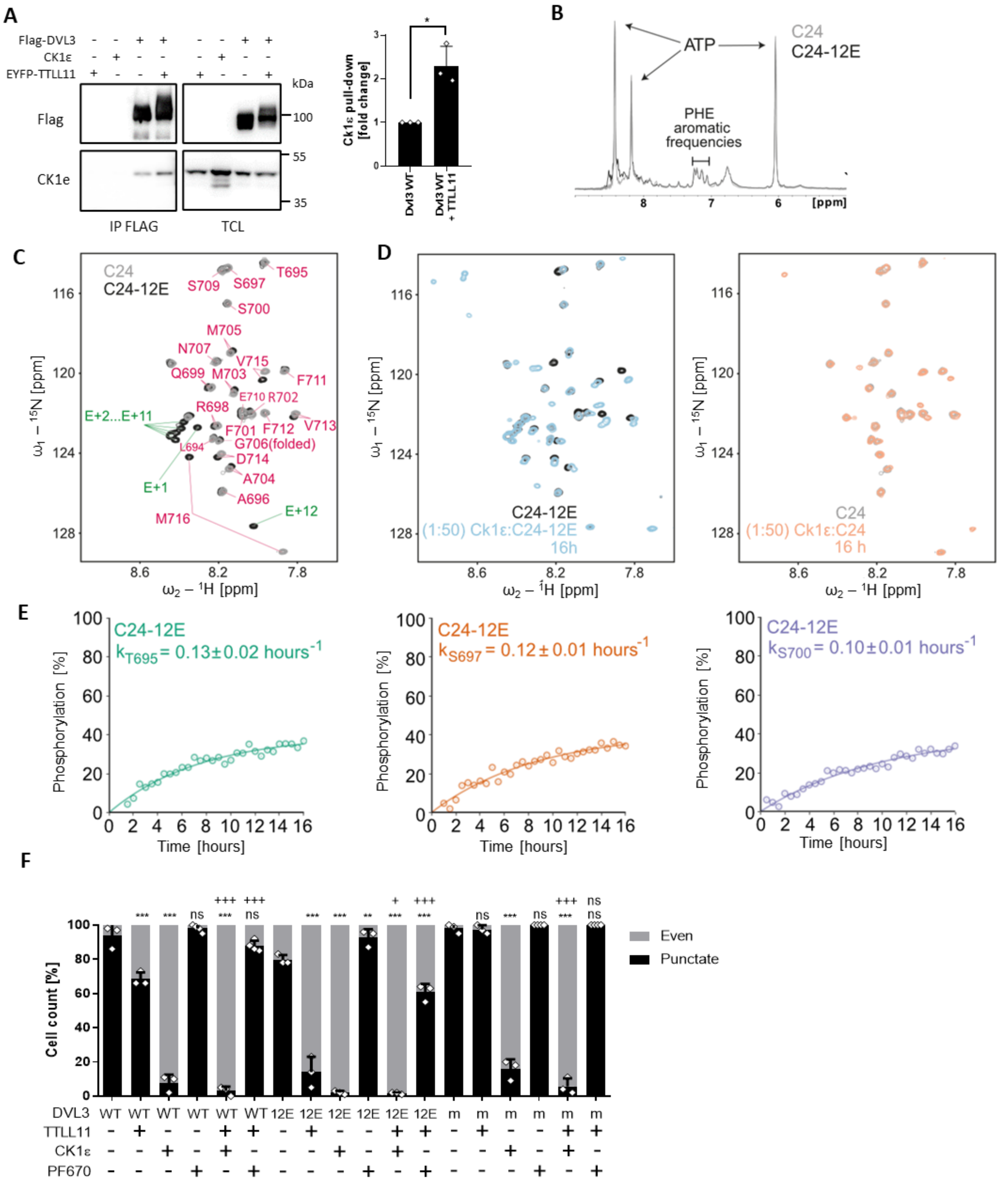
A
Flag-DVL3
CK1ε
EYFP-TTLL11
kDa
Flag
CK1e
100
55
35
IP FLAG
TCL
Ck1ε pull-down [fold change]
Dvl3 WT
Dvl3 WT + TTLL11
B
ATP
C24
C24-12E
PHE aromatic frequencies
[ppm]
C
C24
C24-12E
ω2 – 1H [ppm]
ω1 – 15N [ppm]
D
C24-12E
(1:50) Ck1ε:C24-12E
16h
C24
(1:50) Ck1ε:C24
16 h
E
C24-12E
k T695 = 0.13±0.02 hours-1
k S697 = 0.12±0.01 hours-1
k S700 = 0.10±0.01 hours-1
Phosphorylation [%]
Time [hours]
F
Cell count [%]
Even
Punctate
DVL3
TTLL11
CK1ε
PF670

**Fig. S9. The effect of polyglutamylation on the DVL3 phosphorylation by CK1ε.**

(**A**) Co-immunoprecipitation of overexpressed DVL3 with endogenous CK1ε in presence and absence of co-expressed TTLL11 in HEK293T cells. The cells were transfected as indicated and pull-down was performed using specific antibodies. The graph quantifies the interaction between DVL3 and CK1ε upon co-expression of TTLL11. CK1ε band intensity was normalized to Flag intensity and results are represented as a fold change to CK1ε pull-down by DVL3 WT. Statistical significance was analyzed by one-sample t-test with theoretical mean = 1; n=3. (**B**) For real-time NMR phosphorylation reactions, the concentration of DVL3 C-terminal C24 and C24-12E peptides was matched in the aromatic frequencies of phenylalanine residues in 1D spectra. (**C**) Chemical shift assignment of C24 (grey) and C24-12E (black) peptides. (**D**) Overlay of HSQC spectra of non-phosphorylated form (grey and black for C24 and C24-12E, respectively) and phosphorylated form of the peptide (red and blue for C24 and C24-12E, respectively) after 16 h of the phosphorylation reaction. (**E**) Fraction of phosphorylated T695, S697 and S700 of C24-12E peptide over time. Rate constants were estimated from the mono-exponential fitting to experimental data. (**F**) Analysis of punctate vs. even subcellular localization of overexpressed DVL3 and its polyglutamylation variants (DVL3m = DVL3 E710A/D714A) upon co-expression or inhibition of CK1ε by PF670462 (1μM) in HEK293T cells. DVL3 localization was assessed for at least 200 cells in at least three independent replicates. SD and individual data points (white dots) are indicated. Statistically significant difference from the same DVL variant in the persence of TTLL11 (*) and from the same DVL variant in the presence of TTLL11 (+) is indicated. */+ represents $p < 0.05$; **/++ $p < 0.01$; ***/+++ $p < 0.001$; ns - not significant.

Fig. S10

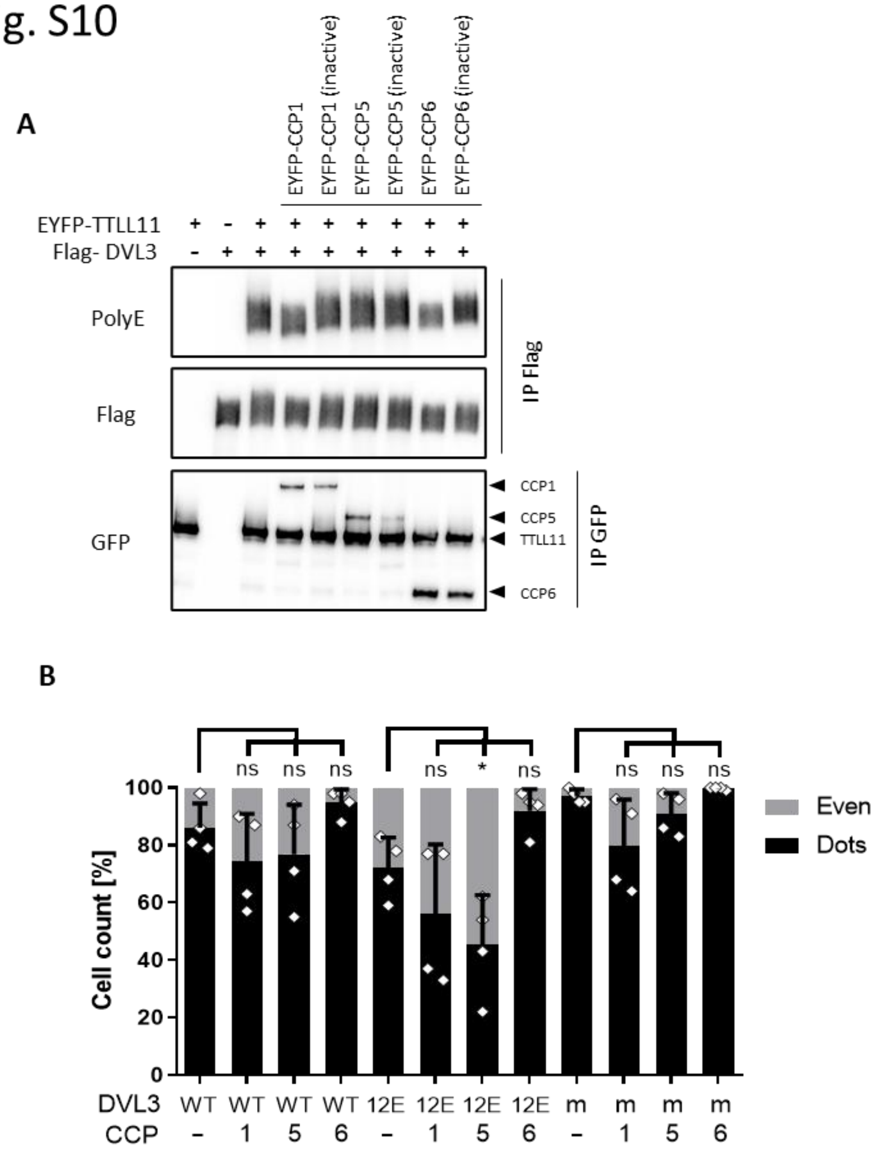


**Fig. S10. Deglutamylation of DVL3 by CCP enzymes.**

(**A**) DVL3 was co-expressed with EYFP-TTLL11 and wild type or inactive EYFP tagged CCP1, CCP5 and CCP6 in HEK293T cells. Upon lysis and immunoprecipitation of DVL3 the level of polyglutamylation was detected using modification-specific antibody PolyE in DVL3 pull-down (IP Flag). The level of CCPs and TTLL11 was monitored in the anti-GFP immunoprecipitates. (**B**) DVL3 variants (DVL3 m stands for E710A/D714A mutant) were co-expressed with EYFP-tagged CCP1, CCP5 and CCP6 in HEK293T cells and subcellular localization of DVL3 was analyzed by immunocytochemistry. Subcellular localization of DVL3 was assessed for at least 200 cells in 3 independent replicates. Mean + SD is shown. Statistical significance in comparison to the condition without CCP for each DVL3 variant is shown. * represents $p < 0.05$; ns - not significant.

#### Table S1.

Identification of TTLL as binding partners of DVL3

| Gene | mW (kDa) | pI (pH) | PLGS Score | Pep-tides | Theore-tical Pep-tides | Cov-erage (%) | Precur-sor RMS Mass Error (ppm) | Pro-ducts | Modi-fied Pep-tides | Pro-ducts RMS Mass Error (ppm) | Pro-ducts RMS RT Error (min) | Pro-tein ID | Identi-fied in | Ref. |
|---|---|---|---|---|---|---|---|---|---|---|---|---|---|---|
| ttll10 | 79 | 9.48 | 96.815 | 1 | 62 | 2.1307 | 7.362 | 8 | 0 | 10.6608 | 0.0386 | 40804 | DVL3 pull-down | (34) |
| ttll3 | 87 | 8.35 | 96.683 | 3 | 61 | 6.0881 | 5.0691 | 17 | 0 | 16.5912 | 0.0399 | 18224 | DVL3 pull-down | (34) |

#### Table S2.

List of plasmids used in this study included in this study.

| Plasmid | Backbone | Insert | Reference |
|---|---|---|---|
| pCDNA3.1-FLAG-mDVL1 | pCDNA3.1 | Flag, mouse DVL1 (1-695) | Kind gift from Madelone Maurice (*80*) |
| pCDNA3.1-FLAG-hDVL2 | pCDNA3.1 | Flag, human DVL2 (1-736) | 3XFlag DVL2 (WT) was a gift from Jeff Wrana. Addgene, # 24802 (*81*) |
| pCDNA3.1-FLAG-hDVL3 | pCDNA3.1 | Flag, human DVL3 (1-716) | Kind gift from Randall Moon (*82*) |
| pCDNA3.1-FLAG-hDVL3 (87-716) | pCDNA3.1 | Flag, human DVL3 (87-716) | Kind gift from Randall Moon (*82*) |
| pCDNA3.1-FLAG-hDVL3 (332-716) | pCDNA3.1 | Flag, human DVL3 (332-716) | Kind gift from Randall Moon (*82*) |
| pCDNA3.1-FLAG-hDVL3 (492-716) | pCDNA3.1 | Flag, human DVL3 (492-716) | Kind gift from Randall Moon (*82*) |
| pCDNA3.1-FLAG-hDVL3 (1-246) | pCDNA3.1 | Flag, human DVL3 (1-246) | Kind gift from Randall Moon (*82*) |
| pCDNA3.1-FLAG-hDVL3 (1-442) | pCDNA3.1 | Flag, human DVL3 (1-442) | Kind gift from Randall Moon (*82*) |
| pCDNA3.1-FLAG-hDVL3 (1-495) | pCDNA3.1 | Flag, human DVL3 (1-495) | Kind gift from Randall Moon (*82*) |
| pDONR221-DVL3 | pDONR221 | human DVL3 (1-716) | DNASU, #HsCD00043699 |
| pDEST Twin-strep-Flag-HALO | pDEDST | Twin-strep,Flag, HALO | Kind gift from Cyril Barinka (*76*) |
| pCDNA3.1-FLAG-hDVL3 E571D/E604D/E693D/E710D | pCDNA3.1 | Flag, human DVL3 (1-716) E571D/E604D/E693D/E710D | This study - mutagenesis of pCDNA3.1-FLAG-hDVL3 |
| pCDNA3.1-FLAG-hDVL3 E710D | pCDNA3.1 | Flag, hDVL3 (1-716) E710D | This study - mutagenesis of pCDNA3.1-FLAG-hDVL3 |
| pCDNA3.1-FLAG-hDVL3 E710A/D714A | pCDNA3.1 | Flag, hDVL3 (1-716) E710A/D714A | This study - mutagenesis of pCDNA3.1-FLAG-hDVL3 |
| pCDNA3.1-FLAG-hDVL3-4E | pCDNA3.1 | Flag, hDVL3 (1-716)-4E | This study - mutagenesis of pCDNA3.1-FLAG-hDVL3 |
| pCDNA3.1-FLAG-hDVL3-4A | pCDNA3.1 | Flag, hDVL3 (1-716)-4A | This study - mutagenesis of pCDNA3.1-FLAG-hDVL3 |
| pCDNA3.1-FLAG-hDVL3 1-709 | pCDNA3.1 | Flag, hDVL3 (1-709) | (*31*) |
| pCDNA3.1-FLAG-hDVL3 1-697 | pCDNA3.1 | Flag, hDVL3 (1-697) | (*31*) |
| pCDNA3.1-FLAG-hDVL3-12E | pCDNA3.1 | Flag, hDVL3 (1-716)-12E | This study - mutagenesis of pCDNA3.1-FLAG-hDVL3 |
| pCDNA3.1-FLAG-hDVL3-E | pCDNA3.1 | Flag, hDVL3 (1-716)-4E | This study - mutagenesis of pCDNA3.1-FLAG-hDVL3 |
| M50 Super8X TopFlash | pTA-Luc | TCF/LEF binding sites | M50 Super 8x TOPFlash was a gift from Randall Moon. Addgene, #12456 (*83*) |

| | | | |
|---|---|---|---|
| pRl-Tk-luc | pRL-TK | luciferase | Promega, #E2241 |
| pcDNA3-hCK1ε wt | pcDNA3 | human CK1ε (1-416) | Kind gift from Lukáš Trantírek (*84*) |
| pCS2+VSVG Lrp6 deltaN | pCS2+ | VSV-G, human Lrp6 (1370–1613) | LRP6 deltaN-pCS2-VSVG was a gift from Xi He. Addgene, # 27281 (42) |
| pCDNA3.1 | pCDNA3.1 | empty vector | Invitrogen, #V79020 |
| pEYFP-spacer-mTTLL1 | pEYFP | mouse TTLL1 (1-423) | Kind gift from Carsten Janke (*3*) |
| pEYFP-spacer-mTTLL2 | pEYFP | mouse TTLL2 (1-540) | Kind gift from Carsten Janke (*3*) |
| pEYFP-spacer-mTTLL3 | pEYFP | mouse TTLL3 (1-927) | Kind gift from Carsten Janke (*3*) |
| pEYFP-spacer-mTTLL4 | pEYFP | mouse TTLL4 (1-1193) | Kind gift from Carsten Janke (*3*) |
| pEYFP-spacer-mTTLL5 | pEYFP | mouse TTLL5 (1-1328) | Kind gift from Carsten Janke (*3*) |
| pEYFP-spacer-mTTLL6 | pEYFP | mouse TTLL6 (1-822) | Kind gift from Carsten Janke (*3*) |
| pEYFP-spacer-mTTLL7 | pEYFP | mouse TTLL7 (1-912) | Kind gift from Carsten Janke (*3*) |
| pEYFP-spacer-mTTLL8 | pEYFP | mouse TTLL8 (1-832) | Kind gift from Carsten Janke (*3*) |
| pEYFP-spacer-mTTLL9 | pEYFP | mouse TTLL9 (1-461) | Kind gift from Carsten Janke (*3*) |
| pEYFP-spacer-mTTLL10 | pEYFP | mouse TTLL10 (1-704) | Kind gift from Carsten Janke (*3*) |
| pEYFP-spacer-mTTLL11 | pEYFP | mouse TTLL11 (1-727) | Kind gift from Carsten Janke (*3*) |
| pEYFP-spacer-mTTLL12 | pEYFP | mouse TTLL12 (1-639) | Kind gift from Carsten Janke (*3*) |
| pEYFP-spacer-mTTLL13 | pEYFP | mouse TTLL13 (1-804) | Kind gift from Carsten Janke (*3*) |
| pEYFP-spacer-hTTLL11 | pEYFP | human TTLL11 (1-800) | Kind gift from Carsten Janke |
| pEYFP-spacer-hTTLL11 (E531G) | pEYFP | human TTLL11 (1-800) E531G | Kind gift from Carsten Janke |
| pcDNA3.1-EYFP-mCCP1 | pcDNA3.1 | mouse CCP1 (1-1218) | Kind gift from Carsten Janke |
| pcDNA3.1-EYFP-mCCP1 (H912S, E915Q) | pcDNA3.1 | mouse CCP1 (1-1218) H912S/E915Q | Kind gift from Carsten Janke |
| pcDNA3.1-EYFP-mCCP5 | pcDNA3.1 | mouse CCP5 (1-886) | Kind gift from Carsten Janke |
| pcDNA3.1-EYFP-mCCP5 (H252S, E255Q) | pcDNA3.1 | mouse CCP5 (1-886) H252S/E255Q | Kind gift from Carsten Janke |
| pcDNA3.1-EYFP-mCCP6 | pcDNA3.1 | mouse CCP6 (1-540) | Kind gift from Carsten Janke |
| pcDNA3.1-EYFP-mCCP6 (H230S, E233Q) | pcDNA3.1 | mouse CCP6 (1-540) H230S/E233Q | Kind gift from Carsten Janke |
| Twin strep-Flag-HALO-DVL3 wt | pDEDST | Twin-strep,Flag, HALO, human DVL3 (1-716) | This study |
| Twin strep-Flag-HALO-DVL3-E | pDEDST | Twin-strep,Flag, HALO, human DVL3 (1-716)-E | This study - mutagenesis of Twin strep-Flag-HALO-DVL3 wt |
| Twin strep-Flag-HALO-DVL3-E71A0/D714A | pDEDST | Twin-strep,Flag, HALO, human DVL3 (1-716) E710A/D714A | This study - mutagenesis of Twin strep-Flag-HALO-DVL3 wt |
| Twin strep-Flag-HALO-ECFP-DVL3-wt | pDEDST | Twin-strep,Flag, ECFP human DVL3 (1-716) | This study |
| Twin strep-Flag-ECFP-DVL3-12E | pDEDST | Twin-strep,Flag, ECFP, human DVL3 (1-716)-12E | This study - mutagenezis of Twin strep-Flag-ECFP-DVL3 |
| Twin strep-Flag-ECFP-DVL3 E71A0/D714A | pDEDST | Twin-strep,Flag, ECFP, human DVL3 (1-716) E71A0/D714A | This study - mutagenezis of Twin strep-Flag-ECFP-DVL3 |
| pET-hDVL3 C24 | pET | $His_6$, ZZ tag, hDVL3 (693-716) | This study |
| pET-hDVL3 C24-12E | pET | $His_6$, ZZ tag, hDVL3 (693-716)-12E | This study |
| pCS2+xDvl3 | pCS2+ | 6xmyc, xDvl3 (1-741) | Kind gift from Alexandra Schambony (*85*) |
| pCS2+xDvl3-4E | pCS2+ | 6xmyc, xDvl3 (1-741)-4E | This study - mutagenezis of pCS2+xDvl3 |
| pCS2+xDvl3-6E | pCS2+ | 6xmyc, xDvl3 (1-741)-6E | This study - mutagenezis of pCS2+xDvl3-4E |

| | | | |
|---|---|---|---|
| pCS2+xDvl3-12E | pCS2+ | 6xmyc, xDvl3 (1-741)-12E | This study - mutagenezis of pCS2+xDvl3-6E |

**Table S3.**

List of transfection conditions in individual experiments.

| Experiment | Plasmid | DNA [ng] | PEI ratio (to final concentration of DNA) | Plate and final concentration of DNA per well [ng] |
|---|---|---|---|---|
| Fig 1A-C; S1A-C | pEYFP-spacer-mTTLL1 | 100 | 4x | 24-well; 500 |
| Fig 1A-C; S1A-C | pEYFP-spacer-mTTLL2 | 300 | 4x | 24-well; 500 |
| Fig 1A-C; S1A-C | pEYFP-spacer-mTTLL3 | 250 | 4x | 24-well; 500 |
| Fig 1A-C; S1A-C | pEYFP-spacer-mTTLL4 | 200 | 4x | 24-well; 500 |
| Fig 1A-C; S1A-C | pEYFP-spacer-mTTLL5 | 500 | 4x | 24-well; 500 |
| Fig 1A-C; S1A-C | pEYFP-spacer-mTTLL6 | 100 | 4x | 24-well; 500 |
| Fig 1A-C; S1A-C | pEYFP-spacer-mTTLL7 | 400 | 4x | 24-well; 500 |
| Fig 1A-C; S1A-C | pEYFP-spacer-mTTLL8 | 400 | 4x | 24-well; 500 |
| Fig 1A-C; S1A-C | pEYFP-spacer-mTTLL9 | 300 | 4x | 24-well; 500 |
| Fig 1A-C; S1A-C | pEYFP-spacer-mTTLL10 | 150 | 4x | 24-well; 500 |
| Fig 1A-C; S1A-C | pEYFP-spacer-mTTLL11 | 150 | 4x | 24-well; 500 |
| Fig 1A-C; S1A-C | pEYFP-spacer-mTTLL12 | 50 | 4x | 24-well; 500 |
| Fig 1A-C; S1A-C | pEYFP-spacer-mTTLL13 | 300 | 4x | 24-well; 500 |
| Fig 1A-C; S1A-C | pEYFP-spacer-hTTLL11 | 150 | 4x | 24-well; 500 |
| Fig 1A-C; S1A-C | pEYFP-spacer-hTTLL11 (E531G) | 150 | 4x | 24-well; 500 |
| Fig 1D; S1D | all | 3000 | 3x | 10cm, 6000 |
| Fig 1E | pEYFP-spacer-hTTLL11 | 3000 | 3x | 10cm, 6000 |
| Fig 2A; S2A | all | 3000 | 3x | 10cm, 6000 |
| Fig S2B | all | 3000 | 3x | 10cm, 6000 |
| Fig 2C-E; S2C-D; S4 | all | 3000 | 3x | 10cm, 6000 |
| Fig 3C; S5A, B | all | 3000 | 3x | 10cm, 6000 |
| Fig 4 | all | 2000 | 3x | 10cm, 6000 |
| Fig S6 B, D | all | 100 | 4x | 24-well; 400 |
| Fig S6C | Super8X TopFlash | 100 | 4x | 24-well; 400 |
| Fig S6C | pRl-Tk-luc | 100 | 4x | 24-well; 400 |
| Fig S6C | pCDNA3.1-FLAG-hDVL3 | 50 | 4x | 24-well; 400 |
| Fig S6C | pCDNA3.1-FLAG-hDVL3-4E | 50 | 4x | 24-well; 400 |
| Fig S6C | pCDNA3.1-FLAG-hDVL3-12E | 50 | 4x | 24-well; 400 |
| Fig S6C | pCDNA3.1-FLAG-hDVL3-E710A/D714A | 50 | 4x | 24-well; 400 |
| Fig S6E | all | 1000 | 4x | 6cm; 2000 |
| Fig 5A,B,D,E; S7A,B | all | 150 | 4x | 24-well; 300 |
| Fig 5C; S7C | all | 100 | 4x | μ-Slide 8 well; 200 |
| Fig S8A; 6B; S9A | all | 3000 | 3x | 10cm, 6000 |
| Fig S8F; 6A,C; S9B | all | 150 | 4x | 24-well; 300 |

**Table S4.**

List of primers used for site-directed mutagenesis.

| Original vector | Primer | Sequence (5'-3') |
|---|---|---|
| pCDNA3.1-FLAG-hDVL3 E710D | DVL3 E571D FWD | ccgactgccatcgctgtgctgactgct |
| pCDNA3.1-FLAG-hDVL3 E710D | DVL3 E571D REV | agcagtcagcacagcgatggcagtcgg |
| pCDNA3.1-FLAG-hDVL3 E710D/E571D | DVL3 E604D FWD | tggtccgaatcgctgccgctgccc |
| pCDNA3.1-FLAG-hDVL3 E710D/E571D | DVL3 E604D REV | gggcagcggcagcgattcggacca |
| pCDNA3.1-FLAG-hDVL3 E710D/E571D/E604D | DVL3 E693D FWD | ctggcggtcagatccgggggcac |
| pCDNA3.1-FLAG-hDVL3 E710D/E571D/E604D | DVL3 E693D REV | gtgcccccggatctgaccgccag |
| pCDNA3.1-FLAG-hDVL3 | DVL3 E710D FWD | catcacatccacaaagaaatcactggggtttcccatg |
| pCDNA3.1-FLAG-hDVL3 | DVL3 E710D REV | catgggaaaccccagtgatttctttgtggatgtgatg |
| pCDNA3.1-FLAG-hDVL3 | Dvl3-E FWD | cgaaaagggcgaattctcattccatcacatccacaaagaac |
| pCDNA3.1-FLAG-hDVL3 | Dvl3-E REV | gttctttgtggatgtgatggaatgagaattcgcccttttcg |
| pCDNA3.1-FLAG-hDVL3 | DVL3-4A FWD | gagttctttgtggatgtgatggccgctgcagcctgagaattcgcccttttcggatc |
| pCDNA3.1-FLAG-hDVL3 | DVL3-4A REV | gatccgaaaagggcgaattctcaggctgcagcggccatcacatccacaaagaact |
| pCDNA3.1-FLAG-hDVL3 | DVL3-4E FWD | gagttctttgtggatgtgatggaggaagaggaatgagaattcgcccttttcggatc |
| pCDNA3.1-FLAG-hDVL3 | DVL3-4E REV | gatccgaaaagggcgaattctcattcctcttcctccatcacatccacaaagaact |
| pCDNA3.1-FLAG-hDVL3-4E | DVL3-4E+4E FWD | gaaaagggcgaattctcattcttcttcttcttcctcttcctccatcacat |
| pCDNA3.1-FLAG-hDVL3-4E | DVL3-4E+4E REV | atgtgatggaggaagaggaagaagaagaagaatgagaattcgcccttttc |
| pCDNA3.1-FLAG-hDVL3-8E | DVL3-8E+4E FWD | gaagaggaagaagaagaagaagaggaggaggagtgagaattcgcccttttcg |
| pCDNA3.1-FLAG-hDVL3-8E | DVL3-8E+4E REV | cgaaaagggcgaattctcactcctcctcctcttcttcttcttcttcctcttc |
| pCDNA3.1-FLAG-hDVL3 | DVL3 E710A/D714A FWD | gcgaattctcacatcacagccacaaagaacgcactggggtttcccatg |
| pCDNA3.1-FLAG-hDVL3 | DVL3 E710A/D714A REV | catgggaaaccccagtgcgttctttgtggctgtgatgtgagaattcgc |
| Twin strep-Flag-HALO-DVL3 wt | D3-E FWD | gtacaagaaagctgggctattccatcacatccacaaagaac |
| Twin strep-Flag-HALO-DVL3 wt | D3-E REV | gttctttgtggatgtgatggaatagcccagctttcttgtac |
| Twin strep-Flag-HALO-DVL3 wt | D3 E710A/D714A FWD | gctgggtcacatcacagccacaaagaacgcactggggtttcccatg |
| Twin strep-Flag-HALO-DVL3 wt | D3 E710A/D714A REV | catgggaaaccccagtgcgttctttgtggctgtgatgtgacccagc |
| Twin strep-Flag-HALO-DVL3 wt | D3-4E FWD | gttctttgtggatgtgatggaggaggaggaatagcccagctttcttg |
| Twin strep-Flag-HALO-DVL3 wt | D3-4E REV | caagaaagctgggctattcctcctcctccatcacatccacaaagaac |
| Twin strep-Flag-HALO-DVL3 wt | D3-4A FWD | gttctttgtggatgtgatggccgcggccgcgtagcccagctttcttg |
| Twin strep-Flag-HALO-DVL3 wt | D3-4A REV | caagaaagctgggctacgcggccgcggccatcacatccacaaagaac |
| Twin strep-Flag-HALO-DVL3 wt | D3 (1-709) FWD | caagaaagctgggctaactggggtttcccatg |

| | | |
|---|---|---|
| Twin strep-Flag-HALO-DVL3 wt | D3 (1-709) REV | catgggaaaccccagttagcccagctttcttg |
| Twin strep-Flag-HALO-DVL3 wt | D3 (1-697) FWD | ggaactgaccgccagctagcccagctttcttg |
| Twin strep-Flag-HALO-DVL3 wt | D3 (1-697) REV | caagaaagctgggctagctggcggtcagttcc |
| Twin strep-Flag-HALO-DVL3-4E | D3-4E+8E FWD | gatggaggaggaggaagaagaagaggaagaggaagaggaatagcccagctttcttg |
| Twin strep-Flag-HALO-DVL3-4E | D3-4E+8E REV | caagaaagctgggctattcctcttcctcttcctcttcttcttcctcctcctccatc |
| Twin strep-Flag-HALO-ECFP-DVL3 wt | ΔHALO FWD | ctggctagcaccatggtgagcaagggcgag |
| Twin strep-Flag-HALO-ECFP-DVL3 wt | ΔHALO REV | ctcgcccttgctcaccatggtgctagccag |
| pCS2+xDvl3 | xDvl3-4E FWD | ctggggtgtttgactttctcgaggaagaggaatagaactatagtgagtcgta |
| pCS2+xDvl3 | xDvl3-4E REV | tacgactcactatagttctattcctcttcctcgagaaagtcaaacaccccag |
| pCS2+xDvl3-4E | xDvl3-6E FWD | gactttctcgaggaagaggaagaggaatagaactatagtgagtcgtat |
| pCS2+xDvl3-4E | xDvl3-6E REV | atacgactcactatagttctattcctcttcctcttcctcgagaaagtc |
| pCS2+xDvl3-6E | xDvl3-12E FWD | gaggaagaggaagaggaagaggaagaggaagaggaatagaactatagtgagtcgtat |
| pCS2+xDvl3-6E | xDvl3-12E REV | atacgactcactatagttctattcctcttcctcttcctcttcctcttcctcttcctc |

#### Table S5.

List of antibodies used in the study.

| Antibodies | Source | Company | Dillution | RRID |
|---|---|---|---|---|
| GFP | Rabbit | Fitzgerald, #20R-GR-011 | *Used for IP | AB_1286217 |
| GFP [3H9] | Rat | Chromotek, #ABIN398304 | WB 1:1000 | AB_10773374 |
| PolyE | Rabbit | Adipogen, #AG-25B-0030-C050 | WB 1:1000 | AB_2490540 |
| Actin beta | Rabbit | Cell signaling, #4970 | WB 1:1000 | AB_2223172 |
| DVL2 | Mouse | Santa Cruz, #8026 | WB 1:1000 | AB_627432 |
| DVL3 | Mouse | Santa Cruz, #8027 | WB 1:1000 | AB_627434 |
| CK1e | Mouse | BD Biosciences, #610445 | WB 1:1000 | AB_397819 |
| FLAG | Rabbit | Sigma-Aldrich, #F7425 | *Used for IP (1ul) | AB_439687 |
| FLAG M2 | Mouse | Sigma-Aldrich, #F1804 | WB 1:2000; ICC 1:1000; *used for IP | AB_262044 |
| myc | Rabbit | Sigma-Aldrich, # C3956 | WB 1:1000 | AB_439680 |
| Alexa Fluor 568 | Donkey - anti mouse | Invitrogen, #A10037 | ICC 1:500 | AB_2534013 |
| Anti-Mouse IgG - HRP | Goat | Sigma-Aldrich, #A4416 | WB Secondary 1:5000 | AB_258167 |
| Anti-Rabbit IgG - HRP | Goat | Sigma-Aldrich, #A6667 | WB Secondary 1:5000 | AB_258307 |
| Anti-Rat IgG - HRP | Goat | Sigma-Aldrich, #A9037 | WB Secondary 1:5000 | AB_258429 |
